# Perceptual Foundation and Extension to Phase Tagging for Rapid Invisible Frequency Tagging (RIFT)

**DOI:** 10.1101/2023.12.22.572966

**Authors:** Eelke Spaak, Floortje G. Bouwkamp, Floris P. de Lange

## Abstract

Recent years have seen the emergence of a visual stimulation protocol called Rapid Invisible Frequency Tagging (RIFT) in cognitive neuroscience. In RIFT experiments, visual stimuli are presented at a rapidly and sinusoidally oscillating luminance, using high refresh rate projection equipment. Such stimuli result in strong steady-state responses in visual cortex, measurable extracranially using EEG or MEG. The high signal-to-noise ratio of these neural signals, combined with the alleged invisibility of the manipulation, make RIFT a potentially promising technique to study the neural basis of visual processing. In this study, we set out to resolve two fundamental, yet still outstanding, issues regarding RIFT; as well as to open up a new avenue for taking RIFT beyond frequency tagging per se. First, we provide robust evidence that RIFT is indeed subjectively undetectable, going beyond previous anecdotal reports. Second, we demonstrate that full-amplitude luminance or contrast manipulation offer the best tagging results. Third and finally, we demonstrate that, in addition to frequency tagging, phase tagging can reliably be used in RIFT studies, opening up new avenues for constructing RIFT experiments. Together, this provides a solid foundation for using RIFT in visual cognitive neuroscience.

## 1. Introduction

The use of periodic visual stimuli to elicit robust stimulus-specific neural activity has a long history in visual cognitive neuroscience (Adrian & Matthews, 1934; Norcia et al., 2015; Vialatte et al., 2010). Typically, individual stimuli, ranging from simple LEDs to complex visual patterns, are flashed to participants at a particular presentation frequency (most commonly between 10 and 30 Hz). This reliably results in a steady-state visual evoked potential or field (SSVEP/SSVEF) at the presentation frequency over occipital regions, measurable by EEG or MEG, respectively. The SSVEP/F is known to have a high signal-to-noise ratio and such ‘tagging’ studies have therefore proven a valuable addition to the visual cognitive neuroscientist’s toolkit (Vialatte et al., 2010).

Recently, driven by technical innovations in presentation equipment, several researchers have pointed out the distinct advantages of presenting rhythmic stimuli not as flashes at relatively low frequency, but as sinusoidally modulated stimuli at higher frequencies. This technique is often referred to as Rapid Invisible Frequency Tagging (RIFT) (Drijvers et al., 2021; Seijdel et al., 2022; Zhigalov et al., 2019). In classical SSVEP/F studies using slower frequencies, the rhythmic modulation is highly apparent to the participants; i.e., they can clearly see the stimuli flicker. Furthermore, rhythmic stimulation at moderate frequencies is known to interact with endogenous neural oscillations in the same frequency range (de Graaf et al., 2013; Haegens, 2020; Keitel et al., 2019; Mathewson et al., 2012; Obleser & Kayser, 2019; Spaak et al., 2014). This poses problems if one wishes to study these endogenous oscillations and their interaction with incoming stimuli (Seijdel et al., 2022). RIFT employs sinusoidally modulated stimuli around 60 Hz, corresponding to somewhat of a ‘sweet spot’ on the frequency axis. This frequency range is high enough that the resulting tagging is thought to be invisible to participants, yet low enough to still result in an appreciable peak in the spectrum of electromagnetic brain activity (resonant amplitude drops with increasing frequency; (Herrmann, 2001; Minarik et al., 2023)). The apparent invisibility allows RIFT to be employed in considerably more naturalistic settings than traditional SSVEP/F (in nature, strong and visible flicker is rare). Furthermore, narrowband modulated stimuli around 60 Hz do not entrain endogenous neural oscillations (Duecker et al., 2021), allowing the study of the latter in stimulated circumstances.

The promises (and existing accomplishments) of RIFT notwithstanding, whether these modulations are truly invisible has so far never been objectively tested. Going forward, RIFT needs a stronger foundation than the subjective, anecdotal, assertion of invisibility by researchers. Therefore, the first goal of the present study is to perform an objective test of the (in)visibility of RIFT. Furthermore, one can apply a sinusoidal modulation of stimulus material in several ways, each of which has a different consequence for the appearance of the stimulus on the screen, and possibly for the neural response as well. As a second objective, we set out to compare several such stimulation protocols. Finally, RIFT is typically focused on tagging different stimuli with different frequencies, as this allows a relatively straightforward analysis of the corresponding brain responses. Given the narrow ‘sweet spot’ of RIFT frequencies, it would be desirable if one could additionally manipulate tagging *phase* and reliably measure the resulting neural response as well. This would considerably increase tagging bandwidth. As a third objective, we therefore test the feasibility of this hitherto unexplored approach.

## 2. Methods

We set out to answer the above questions by means of an experiment in which we presented RIFT-tagged stimuli to participants, tested their behaviour, and recorded MEG data.

### 2.1 Participants

We recruited 14 healthy participants (8 female; 6 male; approximate age 26 ± 6 years (mean ± standard deviation; only year of birth recorded for privacy reasons), range 22–44 years) from the Radboud University participant pool. All participants had normal or corrected-to-normal vision. Participants received € 15 compensation for their participation. The study was approved by the local ethics committee (CMO Arnhem-Nijmegen, Radboud University Medical Center) under the general ethical approval for the Donders Centre for Cognitive Neuroimaging (“Imaging Human Cognition”, CMO 2014/288). Participants provided written informed consent prior to the experiment.

Data from one participant was contaminated by a strong high-frequency drifting artifact due to MEG equipment malfunction. Data from another participant was of poor quality due to excessive head motion. Both these participants are excluded from further analyses, leaving a total of 12 participants on which the reported analyses are based.

### 2.2 Stimuli and task

Participants completed two different tasks: a discrimination task and a passive viewing task. In both tasks, stimuli were circular aperture square wave gratings, presented at a randomly chosen orientation of either -45° or +45°. In passive viewing task blocks, the grating radius was 4 dva (degrees of visual angle), while in discrimination blocks, the radius was 2 dva. Gratings had a small circular central cutout (radius 0.4 dva in passive blocks, 0.2 dva in discrimination blocks) to facilitate fixation. In passive viewing blocks, a fixation dot was presented at all times in the center of the screen.

Each trial began with a fixation dot presented in isolation for a random duration between 500 and 800 ms. In the passive viewing task, this was followed by either one grating presented centrally or two gratings presented, one in either visual hemifield (Figure 1B). Gratings were presented for 1200 ms. Before approximately every 5 trials (interval randomly chosen between 4 and 6 inclusive), the fixation dot would dim, indicating to participants that they had to press a button, in order to keep them alert and fixating. Additionally, participants were instructed that they should blink their eyes during these button press events, to reduce the number of blinks during trials of interest. Data from button press events was not included in any analyses.

**Figure 1:**
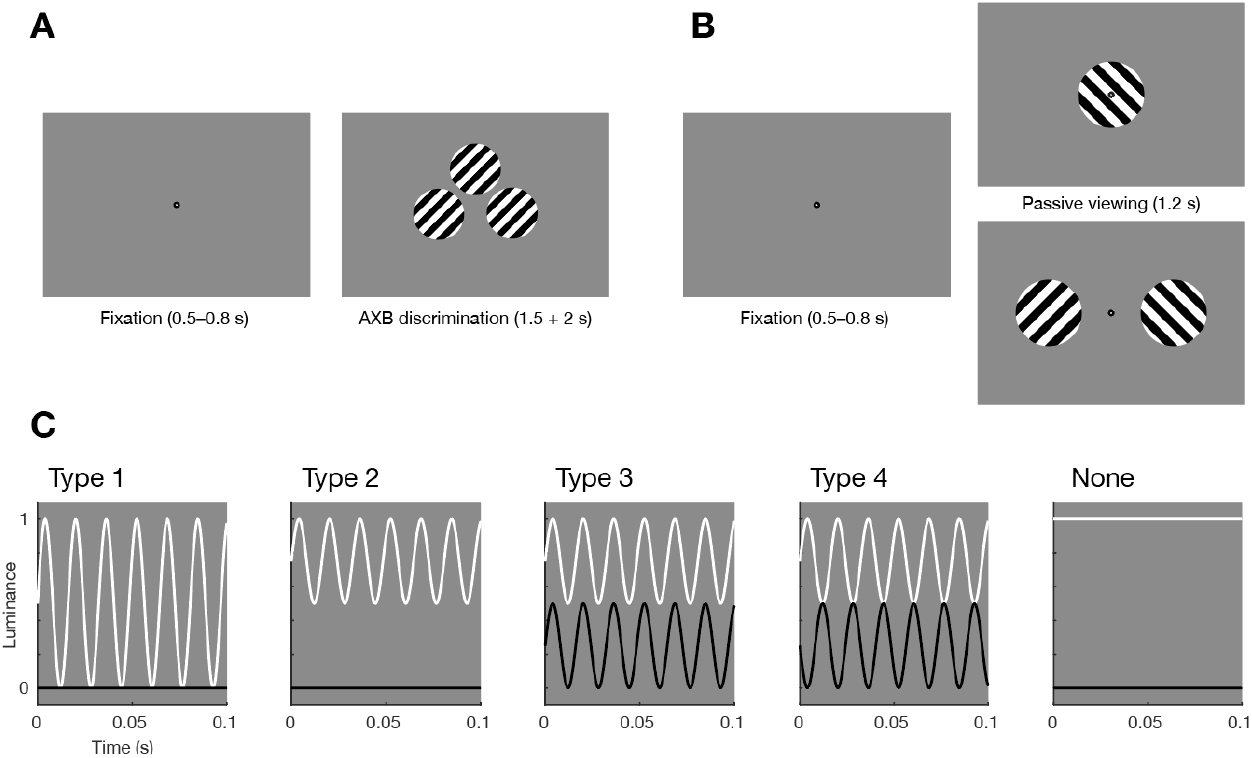
Experimental design and tagging types. **(A)** Setup of the discrimination task. After a brief fixation period, participants were presented with two templates (left and right) and one target (middle top) and had to indicate with a button press whether the target was identical to either the left or right template. **(B)** Setup of the passive viewing task. After a brief fixation period, participants were presented with either a single central stimulus or two stimuli to either side of fixation. An occasional fixation dot dim (not shown) indicated that a button press was required, in order to maintain attention. **(C)** The four different tagging types, illustrated by plotting the luminance over time for the white and black bars of the grating stimuli. See text for details.

In the discrimination task, the fixation dot was followed by three gratings presented on the screen (Figure 1A). The left and right grating served as ‘templates’, the center grating was a ‘target’. Participants were instructed that the center grating was in some way identical to either the left or the right template grating, and their task was to press a left or right button to indicate which of these two templates they believed was identical to the target. Gratings were presented for 1500 ms, and participants had an additional 2000 ms after that during which to provide their response. Such a discrimination task is sometimes referred to as an ‘AXB’ task: the objective is to compare a target (X) to two provided templates (A/B) (Greenaway, 2017). Such a task should be more sensitive to pick up any perception of tagging in participants than a simple two-stimulus ‘pick the tagged one’ task, since at any time participants could compare the target to simultaneously presented reference templates.

### 2.3 Tagging manipulation

We applied RIFT according to four protocols or ‘tagging types’, which we refer to simply as types 1–4 (Figure 1C). Type 1 corresponds to full amplitude luminance tagging: the white bands (i.e., those bands that are white in the untagged grating) of the grating oscillate between 0% and 100% luminance, while the black bands remain at 0% constant. Note that this leads to a time-average luminance of the white bands of 50%, or mid-grey. That means that the ‘white’ bands will become invisible against a typical mid-grey background. This does not pose an issue for the tagging per se, but the stimulus would then appear clearly different from the other tagging type conditions (specifically, only black bars against a grey background). Therefore, to keep appearance more uniform, we presented Type 1 trials always against a full-black (0%) background, while the other tagging types are presented against a mid-grey background. Type 2 tagging corresponds to the white bands oscillating between 50% and 100% luminance (half amplitude luminance tagging), with black bands again at a constant 0% black. This has the advantage that a mid-grey background is again useable, since the ‘white’ bands are now at a time-average 75% grey, clearly distinguishable from the 50% background. Type 3 tagging corresponds to the white bands again oscillating between 50% and 100% luminance, while the black bands oscillate between 0% and 50% luminance in-phase. This protocol is what would naturally result if one constructs an arbitrary monochrome stimulus at 50% dynamic range and applies the tagging as a uniform additive luminance modulation on top. Type 4 tagging corresponds again to the white bands oscillating between 50% and 100% luminance, and the black bands between 0% and 50% luminance, but with the important difference that here the tagging oscillations are in anti-phase. In other words, Type 4 tagging corresponds to *contrast* tagging, rather than luminance tagging. This protocol corresponds to constructing an arbitrary monochrome stimulus at full dynamic range and manipulating its contrast (at full amplitude) over time.

Tagging was applied at 60 Hz, unless specified otherwise (see below). Typically, RIFT manipulation is performed by keeping tagging phase constant across trials, allowing for a straightforward analysis. Since one of the goals of this study was to test whether phase can reliably be reconstructed as well, the passive viewing task included both blocks with a constant phase across trials, as well as blocks where tagging phase was random across trials.

### 2.4 Experimental design

Participants first completed 9 blocks of passive viewing trials, in random order. Tagging type and stimulus condition (one or two stimuli) were manipulated blockwise. Four blocks were single-stimulus (corresponding to tagging types 1–4), fixed phase; two blocks were single-stimulus (tagging types 1 and 4), random phase; two blocks were two-stimulus, with one stimulus at a random phase, and the other at a 90° phase offset. In one block (single stimulus), no tagging was applied at all. Three additional passive viewing blocks were recorded, but not analyzed in the present study: two stimuli tagged at different frequencies (60/66 Hz) for two tagging types (1 and 4), and one block of two stimuli with no tagging. All single-stimulus passive viewing blocks consisted of 50 trials, while two-stimulus blocks consisted of 80 trials.

After the passive viewing blocks, participants engaged in two blocks that included an attentional manipulation (two tagged stimuli on the screen, spatial attention cue). Data from these blocks are not analyzed in the present study. (Any data not analyzed here is still included in the open data release that accompanies this study.)

Finally, participants completed three blocks (50 trials each) of the discrimination task. In two of these (tagging types 1 and 4), one template (left or right) stimulus was always not tagged, while the other was tagged at 60 Hz. In the third discrimination block, both stimuli were tagged (type 1), with one at 60 Hz and the other at 66 Hz. The target stimulus was always randomly chosen to be identical to either the left or right template.

### 2.5 Apparatus

Stimuli were presented using Matlab (The Mathworks, Inc., Natick, Massachussetts, United States) and custom-written scripts using the Psychophysics Toolbox (Brainard, 1997), and back-projected onto a translucent screen (48 × 27 cm) using a ProPixx projector (VPixx Technologies, Saint-Bruno, Québec, Canada) driven at a resolution of 960 × 540 and a refresh rate of 1440 Hz. Participants were seated at a distance of 85 cm from the screen.

Brain activity was recorded using a 275-channel axial gradiometer MEG system (VSM/CTF Systems, Coquitlam, British Columbia, Canada) in a magnetically shielded room. During the experiment, head position was monitored using three fiducial coils (nasion/left ear/right ear). Whenever participants’ head movement exceeded ∼5mm, the experiment was manually paused and the head position was shown to the participant, who would subsequently reposition (Stolk et al., 2013). Eye position and blinks were recorded using an Eyelink 1000 eye tracker (SR Research Ltd., Mississuaga, Ontario, Canada). A tagging signal, if present, was always applied not only to the experimental stimuli, but also to a small square drawn in the bottom left corner of the experiment screen. This square was covered in a light-sensitive diode, and data from this sensor was recorded along with the MEG data. All data were on-line low-pass filtered at 300 Hz and digitized at a sampling rate of 1200 Hz. Immediately prior to the MEG session, participants’ headshape and the location of the three fiducial coils were digitized using a Polhemus 3D tracking device (Polhemus, Colchester, Vermont, United States) in order to facilitate subsequent source analysis.

### 2.6 Discrimination task data analysis

We analyzed the data from the three discrimination task blocks (60 Hz vs no tagging, Type 1; 60 Hz vs no tagging, Type 4; 60 Hz vs 66 Hz, Type 1) separately. We analyzed the single trial data using a Bayesian hierarchical logistic regression model (Bernoulli family, logit link), modelling the single-trial response (left or right) as a function of the actually correct response (left or right), plus intercept. Both the intercept and sensitivity (i.e., the regression coefficient for the actually correct response) were included as subject-random factors. Priors for all parameters were chosen to be weakly informative (i.e., wide spread, with regression coefficients’ priors centered on zero). Modelling and prior construction was done using the Bambi toolbox (Yarkoni & Westfall, 2016), with underlying Markov-chain Monte Carlo (MCMC) estimation done using PyMC (Salvatier et al., 2016). We report evidence for the null or alternative hypothesis as Bayes factors, BF_01_ (in favour of null) or BF_10_ (in favour of alternative). Trials with no response due to timeout were excluded from all analyses (2.3% of trials, median across participants). For graphical purposes (Figure 2A), we additionally compute simple accuracy as the proportion of correct responses per participant. Before computing accuracy, we removed data from one participant, in one condition, since they only responded on one trial in that condition. These data (i.e., this single trial) is still included in the Bayesian analysis.

**Figure 2:**
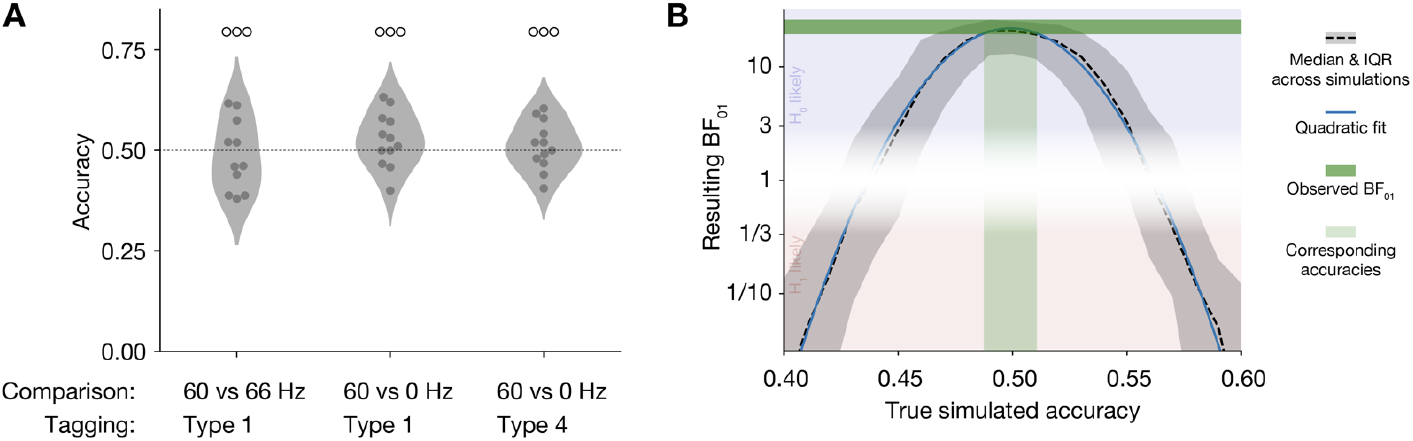
Discrimination task behavioural results. **(A)** Accuracy for the three conditions of the AXB discrimination task. Dots reflect individual participants. Open circles above the density plots reflect strength of evidence for the null hypothesis of random guessing in hierarchical Bayesian logistic regression. ○○○: 10 < BF_01_ < 30. **(B)** Results for sensitivity analysis. BF_01_ are shown for varying true simulated observer accuracy levels. Y-axis is on a log_2_-scale for interpretability. Dashed line shows median across simulations, shading is the inter-quartile range (IQR). Blue line shows a quadratic fit to the median data. Dark green band indicates the minimum and maximum (across the three conditions) observed BF_01_; light green band indicates those (median) accuracies that correspond to these BF_01_.

To put the Bayes factors for the observed data in perspective, we additionally conducted a sensitivity analysis by simulation. For a number of possible true underlying accuracies (from 0.4 to 0.6, inclusive, in steps of 0.01), we simulated a random dataset of identical size (number of participants, number of trials per participant) and spread (standard deviation of observed accuracies across participants) as the observed data. We then subjected those simulated datasets to the same analysis pipeline as above, and recorded the resulting BF_01_. Per accuracy level, we conducted 300 such simulations with a different random initialization.

### 2.7 MEG data preprocessing and trial selection

All MEG preprocessing, spectral analyses, and source modelling were performed using custom-written scripts and the FieldTrip toolbox (Oostenveld et al., 2011). MEG data from the passive viewing blocks were segmented from −0.4 to +1.2 s around each stimulus display onset. Data were screened for outliers including eye blinks or eye movements, MEG SQUID jumps, and muscle artifacts, using a semi-automatic routine (FieldTrip’s ft_rejectvisual), rejecting artifactual segments and/or excessively noisy MEG channels. After artifact rejection, data were downsampled to 600 Hz (after applying an anti-aliasing filter) to speed up subsequent analyses. Infomax Independent Component Analysis (ICA; (Bell & Sejnowski, 1995)) was then used to clean the data of artifacts caused by ongoing cardiac activity and any residual eye movements.

### 2.8 MEG spectral analysis

Fourier spectra for the passive viewing trials were obtained by segmenting the passive viewing trials from 0.2 to 1.2 s after stimulus onset, yielding 1 s-long data segments and a corresponding 1 Hz frequency resolution. A boxcar (flat) taper was used to minimize power smearing across neighbouring frequencies. Fourier spectra were then transformed to power spectra or inter-trial coherence (ITC) using standard methods, or to brain-to-tagging coherence. While ITC (Delorme & Makeig, 2004) measures the consistency of the phase of the MEG data across trials (which we report in Figure 4 and corresponding analyses, for comparison), with brain-to-tagging coherence (labelled ‘phase-corrected’ in Figure 4) we measure the consistency of the phase difference between a corrected tagging signal and the MEG data.

**Figure 3:**
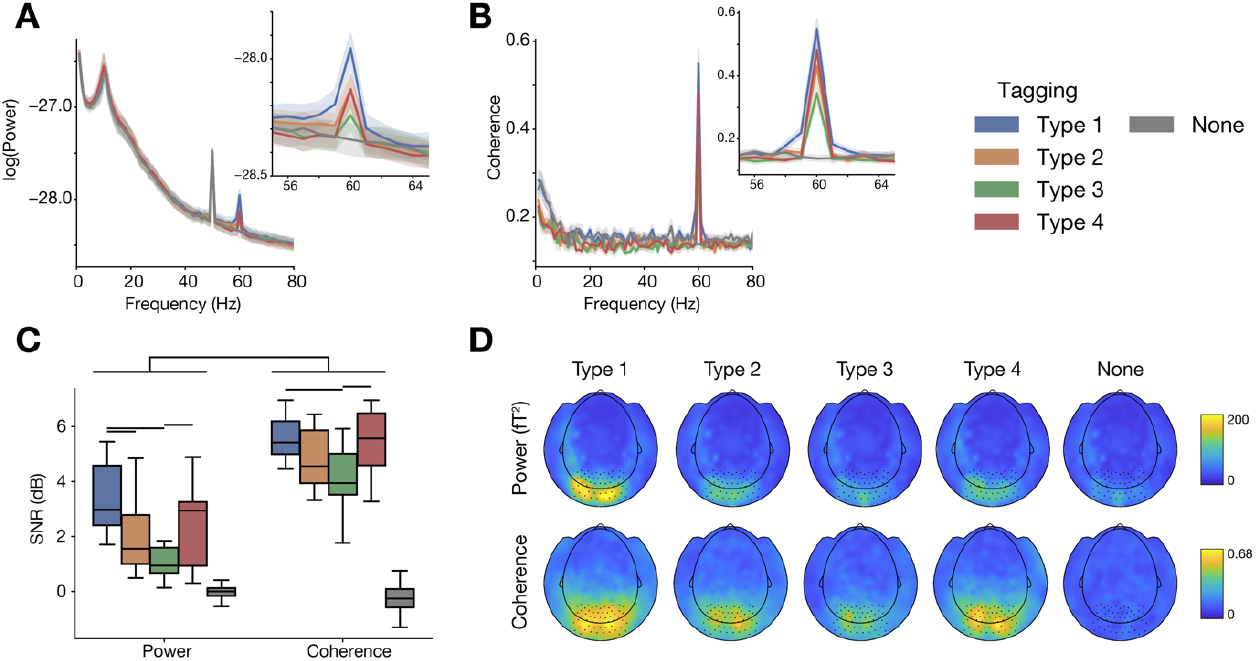
Tagging results for different tagging types (fixed phases, single stimulus on screen). **(A)** Power spectra (log_10_-transformed) for the four tagging types and the no-tagging condition. Noticeable are peaks in the alpha frequency range (∼10 Hz), power line noise (50 Hz) and, importantly, the tagging frequency (60 Hz; only for the tagging conditions). Shading reflects standard error of the mean across participants. Inset shows a zoomed-in version of the spectrum around 60 Hz. Spectra reflect a large set of parieto-occipital sensors, highlighted in panel D. **(B)** Spectra of brain-to-tagging coherence, colours and conventions as in panel A. **(C)** Power and coherence at the frequency of interest (60 Hz), expressed as signal-to-noise ratio (SNR). Box plots reflect the median and inter-quartile range across participants. Horizontal bars indicate differences for which the comparison yields BF_10_ > 3. **(D)** Topography of MEG responses at 60 Hz. Highlighted sensors are those averaged over for the other panels.

**Figure 4:**
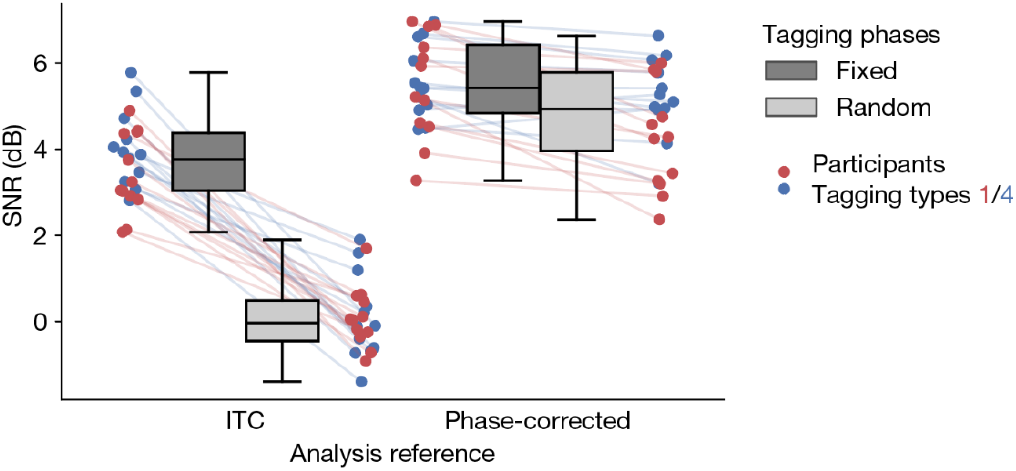
Tagging signal-to-noise ratio (SNR) for fixed and random phase trials, analyzed using inter-trial coherence (ITC) or brain-to-tagging phase corrected coherence.

To obtain this corrected tagging signal, we use: (1) the stored tagging signal as constructed during the experimental session (i.e., the generated sinusoid that was used to tag the stimuli, and also stored to disk), (2) the data from the light sensor attached to the MEG projection screen, and (3) a recording of any missed ‘flips’ (i.e., graphics card buffer updates) during the experimental session (Brainard, 1997). The corrected tagging signal is constructed per trial by first identifying the time point at which the light sensor data ramps up above baseline levels. The stored tagging signal is aligned to that zero point. Then, for any flip that was missed by more than 1 ms, we introduce a matching delay in the corrected tagging signal. (We note that future experiments may employ better (monotonic, ideally linear) analog light sensors which may make this correction procedure redundant (one can then rely on a perfectly measured tagging signal), but the point stands that the stored tagging signal by itself is not optimal for brain-to-tagging coherence.)

To compute signal-to-noise ratio (SNR) from power or coherence spectra, we compared the power or coherence at the frequency of interest (foi) of 60 Hz to the average of neighbouring frequencies up to 5 Hz away, excluding the frequency bins immediately adjacent (i.e., neighbours are [foi – 5, foi – 2] ∪ [foi + 2, foi + 5]), using decibel units:

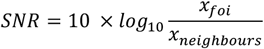

### 2.9 Disentangling orthogonal tagging phases

To analyze the data from the passive viewing task with two stimultaneously presented stimuli, tagged at identical frequencies, but at a relative phase shift of 90°, we adopted the following four-step approach: (1) identify an optimal brain-to-tagging phase lag per participant; (2) project the data onto that phase lag and the corresponding 90° shifted lag; (3) determine the optimal single-dipole pattern explaining the phase-projected data; (4) test for differences between the resulting dipoles (Figure 5A). We elaborate on each of the four steps below.

**Figure 5:**
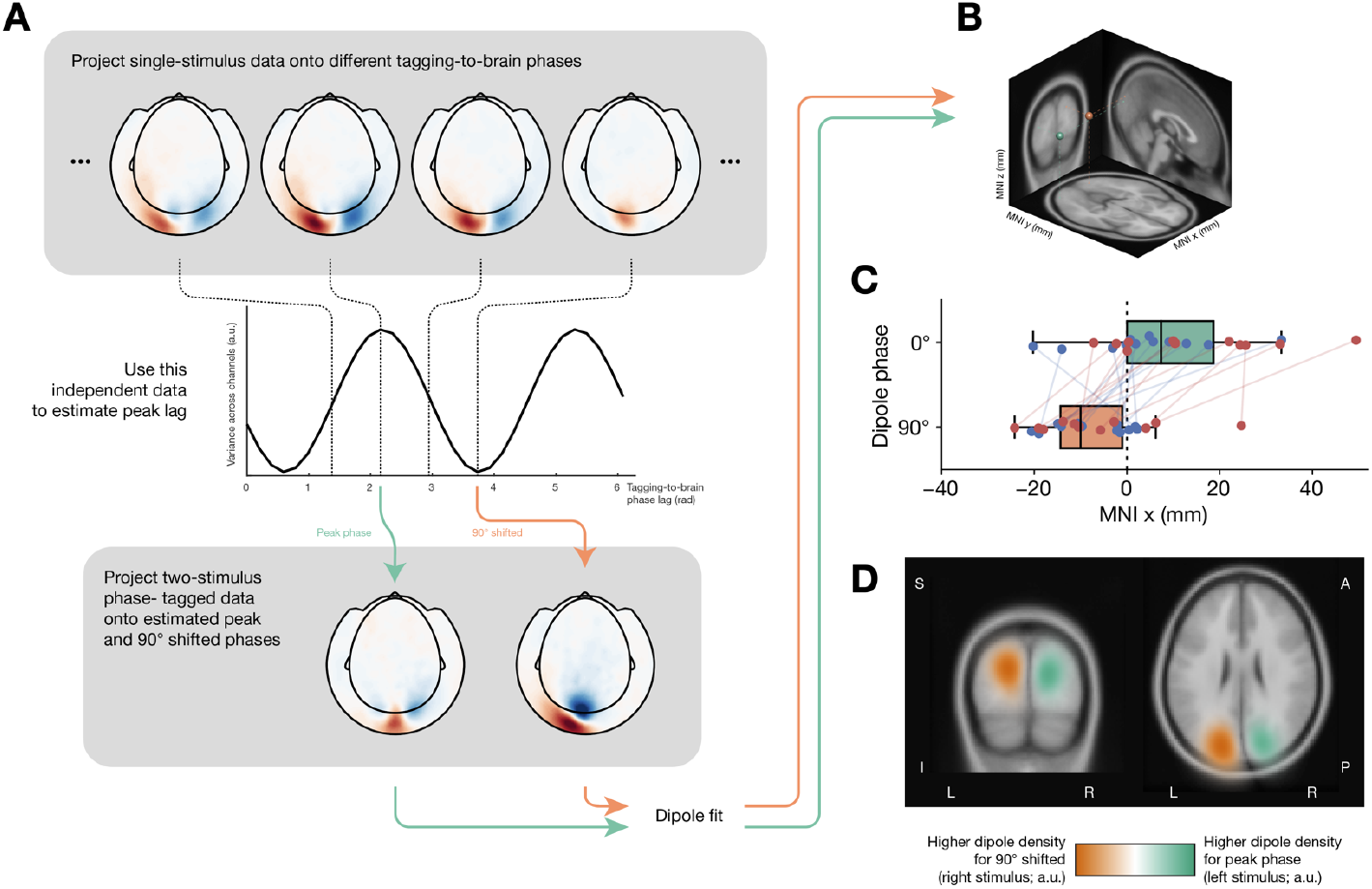
Using RIFT phase tagging. **(A)** Analysis pipeline. Per-channel complex coherency for the single-stimulus (random phase) data is projected onto different phase lags (top). This yields a profile of tagging-to-brain phase lag (middle). The coherency for the two-stimulus data (both stimuli tagged at 60 Hz, always at 90° phase relative to each other) are projected onto both the peak phase lag estimated in this fashion, and the phase 90° away. The resulting projected topographies (bottom) are then subjected to equivalent current dipole (ECD) fitting. **(B)** The two resulting dipoles for a single representative participant. Green corresponds to the fit for the stimulus left on the screen (phase ϕ), orange corresponds to the stimulus right on the screen (phase ϕ + 90°). **(C)** X-coordinates in Montreal Neurological Institute (MNI) space for the ECD fits for all participants. Left stimulus (green) ends up predominantly in the right hemisphere (x > 0), while the right stimulus (orange) ends up predominantly in the left hemisphere (x < 0). Red/blue dots are individual participants, tagging types 1 (red) and 4 (blue). **(D)** Volumetric density plot of all ECD fits, difference between the left and right stimulus, projected onto the template MNI brain. Peaks for the stimuli are clearly shown in contralateral occipital cortex.

(1) We used the single-stimulus, random phase, data to estimate an optimal tagging-to-brain phase lag per participant. We first corrected the MEG data Fourier coefficients for the actual tagging phase (F_corr_ = F_uncorr_ / F_tagging_), to take into account the random stimulation phase used across trials. Then, we projected these complex Fourier coefficients onto several possible phases (0 to 2π in steps of π/16). Different projected phases result in different projection strengths across MEG channels (Figure 5A, top). The amount of total projected MEG variance (across channels; also known as global field power) is a measure of how strongly the brain responds to the stimulus at any particular phase lag. This allows the determination of the optimal tagging-to-brain phase lag per participant (Figure 5A, middle). (2) Then, we applied the same tagging correction procedure to the two-stimulus, random phase, data, and projected the corrected complex Fourier coeffients onto the previously estimated optimal phase, and the relative 90° shifted phase.

(3) The two resulting phase-projected channel topographies (Figure 5A, bottom) were then subjected to equivalent current dipole (ECD) source modelling (Hämäläinen et al., 1993) (Figure 5B), with a single dipole per topography as a reasonable model for a response in early visual cortex (which we confirmed by inspecting the resulting explained variances). Starting location of the fit was always the central calcarine sulcus (MNI x, y, z coordinates (0, -88, -9) mm). Forward models for the source modelling were constructed by nonlinearly warping a template brain (and scalp surface) to the individual participants’ recorded Polhemus headshape data and constructing a single shell volume conduction model (Nolte, 2003). (4) The resulting dipole locations were compared in terms of their left/right displacement (MNI x-coordinates) as a critical test. If the phase tagging were successful, left visual stimuli should result in peaks in right visual cortex, and vice versa.

For illustration (Figure 5D), we furthermore computed a 3D kernel density of resulting dipoles across participants, separately for the 0° (left stimulus) and 90° (right stimulus) cases, using a Gaussian kernel with (isotropic) s.d. 15 mm. We subtracted the two (normalized) densities and projected the difference onto a template MNI brain.

### 2.10 Statistical testing of spectral analysis and source modelling

To assess differences in SNR and ECD position and explained variance, we used hierarchical Bayesian linear models with intercepts as subject-random factors. Models and priors were specified using Bambi (Yarkoni & Westfall, 2016) and MCMC fits were performed using PyMC (Salvatier et al., 2016). Model convergence was verified by inspecting the traces (Kumar et al., 2019) and the Gelman-Rubin statistic 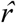.

## 3. Results

### 3.1 Confirming the perceptual invisibility of RIFT

To assess whether participants could observe any difference between tagged and non-tagged stimuli, we adopted an ‘AXB’ paradigm. Participants were presented with two templates (A and B) and a target (X) and had to judge whether the target was identical (“in some way”) to either A or B (Figure 1A). This arguably provides a more sensitive test of subjective perception than merely asking participants to identify which of two stimuli is tagged (or whether a single stimulus is tagged or not) (Greenaway, 2017). There were three comparisons: no tagging versus 60 Hz tagging (Type 1, luminance tagging, see Figure 1C and Methods), no tagging versus 60 Hz tagging (Type 4, contrast tagging), and 60 Hz versus 66 Hz tagging (Type 1). We analyzed these data using hierarchical Bayesian logistic regression, with both the intercepts and sensitivity (i.e., the amount by which the actually correct stimulus influences the response) as subject-random factors. In all three cases, we found strong evidence that participants performed at a level equivalent to random guessing (Figure 2A; 60 vs 66 Hz tagging: BF_01_ = 19.46, 60 Hz vs no tagging (Type 1): BF_01_ = 19.45, 60 Hz vs no tagging (Type 4): BF_01_ = 26.05).

To put the observed evidence for the null hypothesis in perspective, we conducted a sensitivity analysis. For a range of possible true underlying detection accuracies, we simulated random data sets of identical size and spread as the observed data, and subjected these to the identical analysis pipeline as above. Results are shown in Figure 2B (dashed line; blue line indicates quadratic fit). As expected, the evidence for the null hypothesis of guessing (as expressed in BF_01_) peaks when true performance is indeed around chance level (0.5), while a true performance that is clearly above or below chance results in evidence against the null hypothesis (BF_01_ < 1/3). The sensitivity analysis reveals that the BF_01_ for the actually observed data (horizontal green band in Figure 2B) corresponds to a true (median) detection accuracy of between 0.49 and 0.51 (vertical green band). We thereby confirm that the high observed BF_01_ likely corresponds to chance performance.

Finally, we note that at the end of the discrimination blocks, we verbally asked participants whether they noticed anything in particular about the stimuli while they made their decisions. All participants indicated they were entirely guessing; none reported any objective basis for making their left/right decisions. Taking the objective evidence and subjective reports together, we therefore conclude that people indeed cannot see whether RIFT is applied or not.

### 3.2 Full luminance and contrast tagging result in the strongest neural response modulation

We next focused on comparing the different tagging types (Figure 1C; see also Methods) in terms of the resulting neural responses when there is a single tagged (fixed phase) stimulus on the screen which is passively attended (Figure 1B, top). We assessed the neural response modulation both in terms of power and in terms of brain-to-tagging coherence. Power spectra for the different tagging types are shown in Figure 3A. Clearly identifiable are peaks in the alpha range (∼10 Hz), power line noise (50 Hz) and, importantly, at the tagging frequency of 60 Hz. The peak at 60 Hz is absent for the no-tagging condition (most visible in the inset). Coherence spectra (Figure 3B) similarly show a clear peak at 60 Hz for all tagging types.

To make direct comparisons not just among the tagging types, but also between coherence and power, we expressed the tagging as signal-to-noise ratio (SNR) by contrasting the activity at the frequency of interest to neighbouring frequencies. Average results are shown in Figure 3C. Brain-to-tagging coherence analyses provided a considerably higher SNR than power (2.79 ± 0.21 dB difference; posterior mean ± standard deviation; BF_10_ > 10^4^). For power, tagging types 1, 2, and 4 showed demonstrable tagging responses (versus no tagging: BF_10_ > 10^4^), while tagging type 3 resulted in inconclusive evidence (BF_10_ = 1.87). For coherence, all tagging types showed clear tagging responses (all BF_10_ > 10^4^). For both power and coherence, tagging types 1 and 4 resulted in the highest SNR, though with not all pairwise comparisons presenting conclusive evidence (see bars in Figure 3C). Tagging types 1 and 4 resulted in equally high coherence (BF_01_ = 59.54), while their difference was inconclusive for power (BF_10_ = 1.15). Overall, we conclude that brain-to-tagging coherence analysis is preferable over power analysis. Furthermore, tagging types 1 (full luminance tagging) and 4 (contrast tagging) should be preferred over types 2 and 3, when possible. The average sensor topographies (Figure 3D) corroborate this conclusion.

### 3.3 Brain-to-tagging coherence allows the use of random tagging phases

To date, RIFT studies have always employed a fixed tagging phase across trials (Drijvers et al., 2021; Zhigalov et al., 2019). This enables the use of a straightforward coherence-based analysis, namely inter-trial coherence (ITC). One of the goals of the present study was to assess whether phase manipulation could be used for tagging, in addition to frequency. By definition, phase manipulation precludes the use of fixed phases. Therefore, we first tested whether non-constant (i.e., random) tagging phases across trials during the experiment would still allow a robust readout of the tagging signal. We expect the classical ITC analysis to break down here (since it relies on a constant phase). Therefore, we analyzed the data using brain-to-tagging phase corrected coherence, taking the exact tagging phase into account (see Methods for details), and compared the results to ITC. We again focus on the single-stimulus trials here. Given the established equivalence between tagging types 1 and 4, we aggregate across these for the analysis.

There was a clear interaction between tagging phase (fixed/random) and analysis method (ITC/phase-corrected; Figure 4; BF_10_ > 10^4^). As expected, ITC analysis does not detect any tagging signal when phase is randomized across trials (posterior mean SNR ± s.d.: 0.10 ± 0.25 dB, BF_01_ = 54.95), but performs well with constant phases (3.73 ± 0.24 dB, BF_10_ > 10^4^). The evidence for a decrease in ITC with phase randomization was strong (BF_10_ = 42.31). Using phase-corrected brain-to-tagging coherence analysis, tagging is clearly present for both the fixed-phase trials (5.52 ± 0.24 dB, BF_10_ > 10^4^) and the random-phase trials (4.72 ± 0.24 dB, BF_10_ > 10^4^), with no evidence for a difference between the two (BF_10_ = 1.38). From this, we conclude that it is indeed feasible to use a random phase across trials in RIFT studies, and that the neural tagging response can reliably be read out, as long as one takes into account the actual phase that was used in stimulation.

Interestingly, even with a fixed phase across trials, including a phase correction factor substantially increased SNR compared to simple ITC (difference 1.80 ± 0.26 dB, BF_10_ > 10^4^). This is presumably due to subtle but non-constant delays in the experimental apparatus. Measuring the stimulation phase and compensating the tagging reference signal for this allows the recovery of higher SNR, so would in general be recommended, even with constant phases across trials.

### 3.4 Orthogonal phase tagging disentangles multiple same-frequency stimuli

Having established that the use of arbitrary phases across trials for the tagging signal is feasible, we next asked whether phase can be used to uniquely tag different stimuli on the screen. To test this, we presented two stimuli simultaneously to participants (Figure 1B). Both stimuli were tagged at 60 Hz. The phase of the two tagging signals was randomly chosen per trial, with the constraint that the right stimulus always was tagged at a 90° shift relative to the left stimulus.

Since the tagging-to-brain phase lag is different for different participants, properly extracting phase information requires an estimate of this lag. We used the single-stimulus, random phase trials described previously as independent data to perform this estimate (see Methods and Figure 5A for details). Briefly, projecting the single-stimulus data onto various phases allowed an estimation of the peak tagging-to-brain phase lag for each participant. We then projected the two-stimulus data onto that same phase lag, as well as onto the relative 90° shifted phase. By hypothesis, the left stimulus tagging signal should be most strongly visible when projecting at the (independently estimated) peak phase lag, while the right stimulus tagging signal should be dominant when looking at the 90° shifted phase. Taking advantage of the known contralateral projection of visual hemifields, we reasoned that if this hypothesis were true, the non-shifted signal (i.e. the left stimulus) should result in a peak in the right visual cortex, while the 90°-shifted signal (i.e. the right stimulus) should peak in the left visual cortex. We tested this prediction by performing equivalent current dipole (ECD) modelling on the phase-projected channel topographies.

Across participants, dipole fits were of high quality (explained variance r^2^ = 0.78 ± 0.034, posterior mean ± s.d.), with no difference between the peak and 90° shifted phases (Δr^2^ = 0.053 ± 0.037, BF_01_ = 6.46). The resulting dipoles for an example (representative) participant are shown in Figure 5B. Visible are peaks in bilateral occipital cortex, with the left signal peaking in the right hemisphere and vice versa. Critically, also across participants, comparing the MNI x-coordinates for the fits corresponding to the different phase lags revealed that right stimuli peaked in left visual cortex, and vice versa (Δx = 16.39 ± 4.0 mm, BF_10_ = 448.51; Figures 5C/D). We can thus conclude that two stimuli presented simultaneously, tagged with RIFT at an identical frequency but with orthogonal phases, can reliably be disentangled.

## 4. Discussion

We set out to establish a solid empirical foundation for the novel experimental technique of RIFT. First, we confirmed that the ‘I’ in ‘RIFT’ is appropriate: sinusoidally modulated stimuli around 60 Hz are indeed indistinguishable from stimuli that are not modulated at all; the tagging is invisible. Second, we compared four different possible tagging protocols and found that two of them (full-amplitude luminance tagging and contrast tagging) clearly outperform the others. Third, in a direct quantitative comparison, we found that coherence analyses result in higher SNR than power, while power analysis nonetheless still yields an appreciable tagging response. Fourth, we demonstrated that a fixed phase across trials (as typically used) is not a hard requirement for RIFT. Fifth, and finally, we leveraged this potential for non-fixed phases to lay the groundwork for applying phase tagging in future RIFT studies, thereby considerably expanding the potential tagging signal bandwidth.

Concluding subjective invisibility from objective task performance is a notoriously difficult problem in psychophysics (Schmidt, 2015; Vadillo et al., 2022). We believe the conclusion of RIFT’s invisibility is justified here for several reasons. First, we deliberately used an experimental paradigm where a target had to be compared to two templates that were all on the screen simultaneously (an ‘AXB’ task, see (Greenaway, 2017)). This setup eliminates the problematic role of conservative or liberal biases that are present in a more typical one-stimulus setting (where the question may have been “is this flickering or not?”). Second, relatedly, discrimination performance is known to be higher in cases where stimuli are presented simultaneously (Gibson, 1969; Mundy et al., 2009), making the null evidence we found more striking. Third, although we report accuracies in Figure 2, the main evidence for invisibility stems from hierarchical Bayesian logistic regression, in which sensitivity (i.e., the amount by which the actually correct option influences the response) can be well isolated from any remaining bias. Fourth, simulations demonstrated that this analysis pipeline indeed would only reveal very strong evidence in favour of invisibility (as we observed) when actual performance is indistinguishable from chance level. Fifth, and finally, all participants indicated at debriefing that they had no idea on what to base their responses and were entirely guessing. It is noteworthy that the discrimination task was performed at the end of the experimental session, after participants had already experienced RIFT-tagged stimuli for hundreds of trials. Taking all this evidence together, we conclude that RIFT indeed is invisible.

Perhaps unsurprisingly, applying luminance modulation at 100% amplitude (Type 1) resulted in the strongest neural response. A drawback of this type of tagging is that any stimulus that appears white (100%) before applying the modulation is rendered as grey when tagging is applied (time-averaged 50%). Since 50% grey is an often-used background colour in psychophysical experiments, this may not be ideal. Furthermore, how to apply this type of tagging with more complex stimuli than black and white gratings is not well-defined. We examined two flavours of half-amplitude modulation (Types 2 and 3) which may alleviate some of these issues, but found that these resulted in considerably weaker neural tagging responses. Fortunately, we found that contrast modulation (Type 4) produces a tagging response that is equally strong as full-amplitude luminance modulation, while lending itself well to more diverse stimuli. We would recommend future studies to use contrast-modulated tagging in general. Only in specific cases, where stimuli happen to lend themselves well to full-amplitude luminance tagging, would we recommend the use of that tagging type.

The majority of RIFT studies to date has relied on some form of coherence analysis to extract the tagging responses (Drijvers et al., 2021; Zhigalov et al., 2019). By computing signal-to-noise ratios (SNR), we were able to directly compare coherence and power estimates of tagging. This way, we confirmed that, indeed, coherence is a more sensitive measure than power. However, it is worth noting that power analyses still resulted in clear tagging responses. Power analyses are more flexible than coherence; for example, they are easily applied at the single-trial level, unlike coherence. Therefore, when such flexibility is required, it is feasible to resort to power analysis (see, e.g., (Bouwkamp et al., 2023)).

Past RIFT studies have relied on a constant stimulation phase across trials to facilitate subsequent analysis, either through inter-trial coherence (ITC; also known as phase-locking factor; (Delorme & Makeig, 2004)) or through computing the power spectrum of the event-related field (ERF). While this indeed makes analyses straightforward, it limits the potential bandwidth by which one can apply RIFT; specifically, it precludes using phase as a tagging information channel. We confirmed that, indeed, these typical analyses break down when phase is non-constant. Importantly, we demonstrated that a relatively straightforward extension of the fixed-phase analysis method (which we call ‘phase-corrected coherence’ or ‘tagging-to-brain coherence’) is able to recover strong neural tagging signals, even in the case of non-constant phases. Interestingly, this technique resulted in higher SNR than ITC even with constant tagging phases, possibly due to non-constant delays in the presentation equipment, leading us to recommend its use in general.

One clear use case for non-constant phases in RIFT is to tag multiple stimuli at the same frequency, but with different phases. Since there is a narrow ‘sweet spot’ on the frequency axis for RIFT (high enough to be invisible, but low enough to result in good neural SNR), such a doubling of tagging bandwidth can be highly desirable. Relying on our discovery that random phases are feasible, we next demonstrated that, indeed, two stimuli simultaneously presented and tagged at the same frequency, but at orthogonal phases, can reliably be disentangled in the neural response. It would be interesting to see future work examine the feasibility of exploiting the phase axis even further, beyond two orthogonal phases. Furthermore, our current approach relied on using independent data to estimate an individual’s tagging-to-brain phase lag before disentangling the neural responses to two stimuli. The amount of data required for such an estimate is low (just a few minutes should be sufficient), so the extra experimental burden for researchers wishing to go down this route is light. Nonetheless, future work may explore different avenues for estimating the optimal phase lag, possibly without the use of independent (in essence ‘phase localizer’) data.

Previous authors have emphasized the advantages of RIFT, like its leaving endogenous neural oscillations unperturbed, and of course its (hitherto alleged) invisibility (Seijdel et al., 2022). Invisible flicker by itself is already much more naturalistic than classical SSVEP/F paradigms, with highly apparent luminance modulation. We speculate that RIFT may offer a unique way forward in naturalistic visual cognitive neuroscience. The complex stimuli inherent in the natural visual world, and therefore those used in naturalistic experiments, do not lend themselves as easily to the rapid ‘readout’ of neural responses in M/EEG (whether via univariate or multivariate analyses) as well-controlled experimental stimuli (Sonkusare et al., 2019). There is often simply too much going on to be able to reliably say that this particular neural response was a consequence of that visual stimulus. With RIFT, one can tag only certain elements of the visual scene (or movie), without reducing the level of naturalism, and still be able to reliably identify the neural response corresponding to only specific spatial locations or objects. This is a considerable and unique advantage. Of course, further methodological work is needed before RIFT’s potential can be leveraged in naturalistic settings (e.g., do we see strong response modulation also for tagged objects within natural scenes?). Nonetheless, we expect to see a happy marriage between RIFT and naturalistic visual cognitive neuroscience. With such a promising future ahead, RIFT does need a solid empirical foundation, and we hope to have provided one in this paper.

## Data and Code Availability

[All data (with the exception of privacy-sensitive headshape data) and analysis code will be made fully publicly available, under an open license, upon acceptance. Sharing will be done via the Donders Repository (data, persistent DOI) and GitHub (code).]

## Author Contributions

Eelke Spaak: Conceptualization, Methodology, Software, Investigation, Writing – Original Draft, Visualization, Supervision.

Floortje G. Bouwkamp: Conceptualization, Investigation.

Floris P. de Lange: Conceptualization, Writing – Review & Editing, Funding Acquisition.

## Funding

This work was partially funded by the EC Horizon 2020 program (ERC starting grant 678286 awarded to FPdL).

## Declaration of Competing Interests

We declare no competing interests.

## Acknowledgements

We would like to thank Virág Fodor for help in data acquisition.

## Notes

### Competing Interest Statement

The authors have declared no competing interest.

